# Epigenetic and genetic differentiation between *Coregonus* species pairs

**DOI:** 10.1101/2022.11.18.517126

**Authors:** Clare J Venney, Claire Mérot, Eric Normandeau, Clément Rougeux, Martin Laporte, Louis Bernatchez

**Affiliations:** Institut de Biologie Intégrative et des Systèmes (IBIS), Université Laval, Québec, Canada; UMR 6553 Ecobio, OSUR, CNRS, Université de Rennes, Rennes, France; Ministère des Forêts, de la Faune et des Parcs (MFFP), Québec, Québec

**Keywords:** DNA methylation, speciation, whitefish, mutagenesis, genetic divergence, genetic assimilation

## Abstract

Phenotypic diversification is classically associated with genetic differentiation and gene expression variation. However, increasing evidence suggests that DNA methylation is involved in evolutionary processes due to its phenotypic and transcriptional effects. Methylation can increase mutagenesis and could lead to increased genetic divergence between populations experiencing different environmental conditions for many generations, though there has been minimal empirical research on epigenetically induced mutagenesis in diversification and speciation. Whitefish, freshwater members of the salmonid family, are excellent systems to study phenotypic diversification and speciation due to the repeated divergence of benthic-limnetic species pairs serving as natural replicates. Here we investigate whole genome genetic and epigenetic differentiation between sympatric benthic-limnetic species pairs in lake and European whitefish (*Coregonus clupeaformis* and *C. lavaretus*) from four lakes (N=64). We found considerable, albeit variable, genetic and epigenetic differences between species pairs. All SNP types were enriched at CpG sites supporting the mutagenic nature of DNA methylation, though C>T SNPs were most common. We also found an enrichment of overlaps between outlier SNPs with the 5% highest F_ST_ between species and differentially methylated loci. This could possibly represent differentially methylated sites that have caused divergent genetic mutations between species, or divergent selection leading to both genetic and epigenetic variation at these sites. Our results support the hypothesis that DNA methylation contributes to phenotypic divergence and mutagenesis during whitefish speciation.

**Significance statement:** DNA methylation is an epigenetic mark known to change in response to the environment and induce genetic modifications such as point mutations, though its implications for evolution and speciation have not been thoroughly studied. We find considerable but variable genetic and epigenetic variation between whitefish benthic-limnetic species pairs, highlighting the potential for DNA methylation to contribute to mutagenesis and genetic evolution. Our study provides evidence that DNA methylation could have contributed to whitefish speciation, both through initially plastic methylation changes and by driving genetic divergence between species pairs.

## Introduction

Speciation has long been a focus of evolutionary biology, with recent research expanding into the role of genomics in reproductive isolation, phenotypic diversification, and species divergence (Seehausen et al. 2014; Marques et al. 2019). Speciation can occur rapidly despite slow mutation rates, sometimes due to new combinations of standing genetic variation (Marques et al. 2019). Phenotypic plasticity can also promote speciation, particularly when allopatric populations acclimate to their respective environments, phenotypes are partially genetically controlled, and capacity for plasticity is lost over time (Pfennig et al. 2010). Phenotypic changes can occur through altered gene expression (Whitehead & Crawford 2006) which can lead to phenotypic divergence and speciation, as documented between Arctic charr ecotypes (*Salvelinus alpinus*; Gudbrandsson et al. 2018; Jacobs and Elmer 2021), between Chinook and Coho salmon and their hybrids (*Oncorhynchus tshawytscha* and *O. kisutch*; Mckenzie et al. 2021), and between sympatric lake whitefish species (*Coregonus* sp.; Derome et al. 2006; St-Cyr et al. 2008; Rougeux et al. 2019b). Transcriptomic variation associated with lip size has been identified in the Midas cichlid species complex (*Amphilophus citrinellus;* Manousaki et al. 2013) and sequence variation in transcribed regions has been reported between *A. astorquii* and *A. zaliosus* (Elmer et al. 2010). Transcriptomic differences generally increase with taxonomic distance for closely related species (Whitehead & Crawford 2006) and may contribute to phenotypic differences among species (Whitehead & Crawford 2006; Pavey et al. 2010).

Therefore, the mechanisms underlying adaptive phenotypic differences during speciation likely involve both phenotypic plasticity and genetic variants. There has been increasing interest in the role of DNA methylation as a plastic mechanism contributing to speciation (Vogt 2017; Ashe et al. 2021). DNA methylation is the addition of a methyl group to the cytosine of CpG sites in vertebrates (a cytosine followed by a guanine in the DNA sequence), often resulting in altered transcription without a change in DNA sequence (Bird 2002). DNA methylation is sensitive to the environment and epigenetic marks are known to be affected by several factors such as temperature (Ryu et al. 2020; Beemelmanns et al. 2021; McCaw et al. 2020; Venney et al. 2022), salinity (Hu et al. 2021; Artemov et al. 2017; Heckwolf et al. 2020), and rearing environment (Leitwein et al. 2021; Le Luyer et al. 2017; Wellband et al. 2021; Berbel-Filho et al. 2020; Venney et al. 2020). DNA methylation can alter phenotype (Anastasiadi et al. 2021; Vogt 2021), with evidence for phenotypically-linked epigenetic divergence associated with spawning tactics in capelin (*Mallotus villosus*; (Venney et al. 2023) and between four sympatric Arctic charr morphs (Matlosz et al. 2022). Due to the potential role of DNA methylation in plasticity and phenotypic diversification, it may contribute to speciation (Vogt 2017; Ashe et al. 2021; Stajic & Jansen 2021; Laporte et al. 2019). A recent simulation study showed that epigenetic plasticity can promote speciation when it reduces the fitness of migrants and hybrids but can prevent genetic adaptation and speciation if epigenetic adaptation occurs, precluding the need for genetic adaptation (Greenspoon et al. 2022). Empirical evidence for the role of DNA methylation in reproductive isolation and speciation is also emerging (Laporte et al. 2019). Methylation differences detected through methylation-sensitive amplified polymorphism, but not genetic differences, were predictive of behavioural isolation between 16 species of darters (*Ulocentra*, *Nanostoma,* and *Etheostoma*; Smith et al. 2016). Another study showed considerable epigenetic differences between six phenotypically divergent species of Lake Malawi cichlids (Vernaz et al. 2022). DNA methylation can affect transcription (Li et al. 2019) and phenotype (Anastasiadi et al. 2021; Vogt 2021) and could result in initial plastic phenotypic responses that lead to phenotypic diversification and speciation over generations.

DNA methylation is also mutagenic and can generate polymorphism. This is partially due to the spontaneous hydrolytic deamination of methylated cytosine to uracil which is rapidly converted to a thymine tautomer if not corrected by DNA repair enzymes (Gorelick 2003). Spontaneous deamination is ∼3.5 times more likely to occur at methylated cytosines than unmethylated ones (Jones et al. 1992; Gorelick 2003). Different enzymes are involved in base repair of methylated vs unmethylated cytosines, leading to methylated cytosines having mutation rates ∼20 000 times higher than unmethylated ones after accounting for DNA repair efficiency (Gorelick 2003). It is estimated that ∼8 deamination events occur per day in the ∼6 billion bp diploid human genome (Jones et al. 1992) providing a consequential source of novel mutations, particularly in larger genomes. C>T transitions are the most common and explainable epigenetically induced mutation, though there is also evidence for increased C>A and C>G mutations at CpG sites due to mutagen exposure (Tomkova & Schuster-Böckler 2018). On the other hand, DNA methylation can also shield sites from mutagenesis depending on the stimulus or trigger, though the intricacies of when mutagenesis is favoured or prevented remain unclear (Tomkova & Schuster-Böckler 2018). A study in human cell lines (*Homo sapiens*) showed that cytosines with intermediate levels of methylation (20-60%) had the highest mutation rate, even compared to fully methylated sites (Xia et al. 2012). The shielding properties of DNA methylation could also be due to the inability to discern DNA methylation from other methylation-related marks. In particular, 5-methylcytosine can be converted to 5-hydroxymethylcytosine during demethylation (Li et al. 2019). The two cannot be differentiated through bisulfite sequencing (Li et al. 2019), though 5-hydroxymethylcytosine may protect CpG sites from mutation (Tomkova & Schuster-Böckler 2018). It is also possible that higher nucleotide diversity at CpG sites is associated with greater methylation variation at those sites, suggesting weaker selective constraint at those sites (Ord et al. 2023). Conversely, highly methylated sites may be under greater selection to maintain consistently high methylation levels (Ord et al. 2023). Therefore, DNA methylation can not only lead to transcriptional and phenotypic changes but may also influence mutation rates. DNA methylation has thus been proposed as a mechanism for genetic assimilation of phenotypes, i.e., when an environmentally induced phenotype becomes stable and genetically encoded, even in the absence of the original stimulus (Danchin et al. 2019; Nishikawa & Kinjo 2018). However, empirical evidence for methylated sites inducing point mutations and contributing to evolution remains sparse.

The two whitefish sister taxa, lake whitefish (*Coregonus clupeaformis*) in North America and the European whitefish species complex (*C. lavaretus*) in Europe, are well-characterized and relevant systems to study phenotypic diversification and speciation. Lake and European whitefish have evolved separately since they became geographically isolated ∼500 000 years ago (Bernatchez & Dodson 1994, 1991; Jacobsen et al. 2012). Both show repeated independent divergence of sympatric species complexes consisting of a putatively ancestral benthic and derived limnetic species originating from different glacial refugia during the last glaciation period (Østbye, Bernatchez, et al. 2005; Bernatchez et al. 2010; Bernatchez & Dodson 1991; Rougeux et al. 2017; Pigeon et al. 1997) which came into secondary contact ∼12 000 years ago when colonizing postglacial lakes (Rougeux, Gagnaire, Praebel, et al. 2019; Rougeux et al. 2017). In North America, the derived limnetic species evolved to colonize the limnetic zone, leading to differences in diet (Bernatchez et al. 1999), reduced size (Bernatchez et al. 1999), slower growth (Trudel et al. 2001), more slender body morphology (Laporte et al. 2015, 2016), higher metabolic rate (Trudel et al. 2001; Dalziel et al. 2015), and more active swimming behaviour (Rogers et al. 2002). Benthic and limnetic species often coexist in Europe, though some Fenno-Scandinavian and alpine lakes contain up to six sympatric whitefish species (De-Kayne et al. 2022; Østbye, Næsje, et al. 2005). In both North America and Europe, sympatric whitefish are generally reproductively isolated with variable amounts of gene flow depending on the lake (Rougeux et al. 2017; Rogers & Bernatchez 2006) which translates into genetic differentiation (Mérot et al. 2023; Rougeux, Gagnaire, Praebel, et al. 2019; Rogers & Bernatchez 2007; Bernatchez et al. 2010; Dion-Côté et al. 2017; Rougeux et al. 2017; Østbye, Næsje, et al. 2005; De-Kayne et al. 2022; Siwertsson et al. 2013; Østbye et al. 2006), differential transposable element methylation (Laporte et al. 2019), and transcriptional differences (Rougeux, Gagnaire, Praebel, et al. 2019; Derome et al. 2006; Jeukens et al. 2009). Benthic-limnetic species pairs in different lakes thus present a naturally replicated system in which to study the molecular mechanisms associated with speciation.

We assessed the role of genomic and epigenomic variation in whitefish speciation by performing whole genome sequencing (WGS) and whole genome bisulfite sequencing (WGBS) for four benthic-limnetic species pairs (N=64 individuals sequenced in both datasets): lake whitefish from Cliff and Indian Lake, Maine, USA, and European whitefish from Langfjordvatn Lake, Norway and Zurich Lake, Switzerland (Figure 1). The objectives of this study were to (i) determine the extent of whole genome genetic divergence between benthic-limnetic whitefish species pairs, (ii) measure the level of polymorphism at CpG sites relative to the rest of the genome, (iii) characterize whole genome differences in DNA methylation among limnetic-benthic whitefish species pairs, and (iv) assess whether there was an enrichment of outlier SNPs at differentially methylated loci (DMLs), which would support the hypothesis that epigenetically influenced mutagenesis may be creating genetic variation and be involved in speciation. As such, our results provide novel support for the role of DNA methylation in interspecies variation, driving mutagenesis, and genetic evolution between species.

**Fig. 1:**
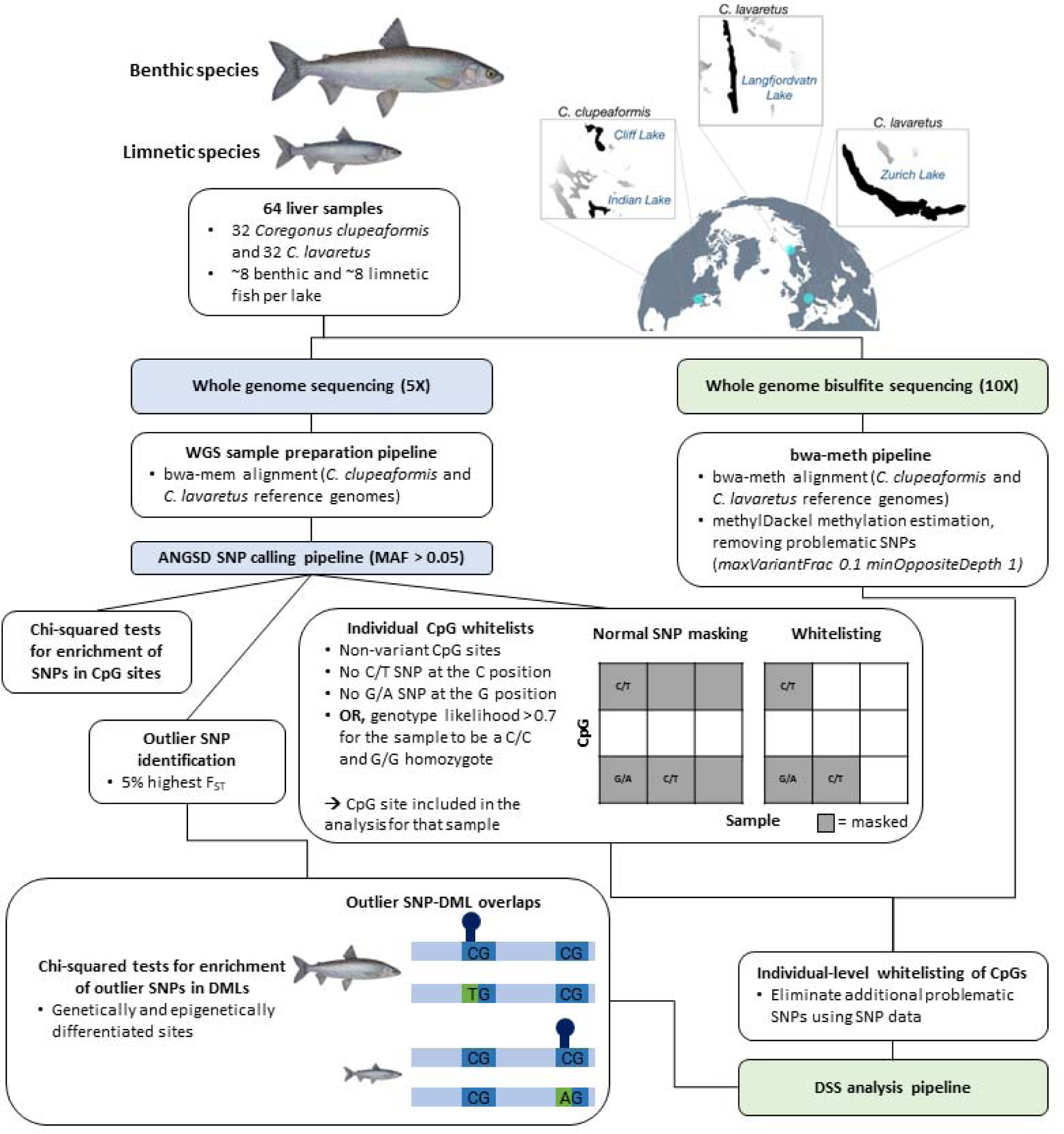
Overview of the sampling design and analysis for sympatric benthic-limnetic species pairs sampled from four lakes. Lake whitefish (*C. clupeaformis*) were sampled from two lakes in North America (Cliff Lake and Indian Lake in Maine, USA) and European whitefish (*C. lavaretus*) were sampled from two lakes in Europe (Langfjordvatn Lake, Norway and Zurich Lake, Switzerland). The analysis pipelines for whole genome sequencing and whole genome bisulfite sequencing are outlined in the flowchart. C/T and G/A SNPs that affect methylation calling are filtered (1) during methylation calling based on settings in methylDackel, and (2) through our whitelisting approach which uses the SNP data for an additional layer of stringent filtration. The *C. clupeaformis* samples were previously analysed in Mérot et al. (2022).

## Results

### Genetic divergence between benthic-limnetic species pairs

The extent of genetic differentiation between benthic and limnetic species varied among lakes. In North America, moderate to high divergence between lakes both within and between species was observed, with greater genome-wide F_ST_ between species in Cliff Lake than Indian Lake (Table 1A). In Europe, F_ST_ estimates for Langfjordvatn Lake and Zurich Lake showed greater differentiation between lakes than between species (Table 1B). In all lakes, genetic differentiation is widespread along the genome (Figure S1). Genetic differentiation between benthic-limnetic species pairs have been comprehensively covered in earlier studies (Gagnaire et al. 2013; Rougeux, Gagnaire & Bernatchez 2019; Rougeux, Gagnaire, Praebel, et al. 2019; Mérot et al. 2023).

**Table 1:**
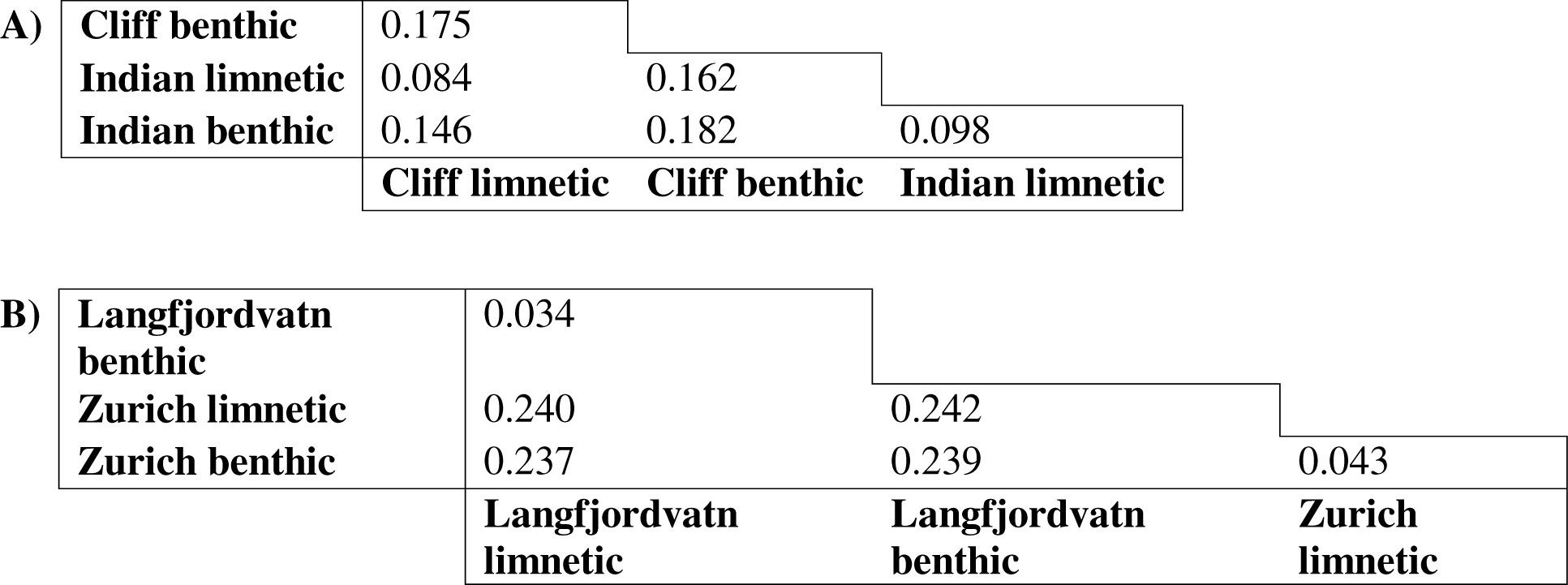
Pairwise F_ST_ matrices for (A) *C. clupeaformis* and (B) *C. lavaretus* reveal variable but considerable genetic divergence between lakes and species.

|  |  |  |  |  |
| --- | --- | --- | --- | --- |
| A) | **Cliff benthic** | 0.175 |  |  |
|  | **Indian limnetic** | 0.084 | 0.162 |  |
|  | **Indian benthic** | 0.146 | 0.182 | 0.098 |
|  |  | **Cliff limnetic** | **Cliff benthic** | **Indian limnetic** |
| B) | **Langfjordvatn benthic** | 0.034 |  |  |
|  | **Zurich limnetic** | 0.240 | 0.242 |  |
|  | **Zurich benthic** | 0.237 | 0.239 | 0.043 |
|  |  | **Langfjordvatn limnetic** | **Langfjordvatn benthic** | **Zurich limnetic** |

### Elevated rate of polymorphism in CpG sites

We analyzed from 11 861 765 to 30 919 358 SNPs per lake at approximately 4X coverage per sample (Table 2) to assess whether SNPs were enriched in CpG sites relative to the rest of the genome to investigate the prediction that CpG sites are mutagenic. We also tested whether the enrichment was specific to (i) C/T and G/A SNPs (to account for the G position of CpG sites), (ii) C/G and G/C SNPs, and (iii) C/A and G/T SNPs. Permutation tests showed that SNPs were significantly enriched in CpG sites for all four lakes (p < 0.0001; Table 2), with none of the 10 000 permutations per test reaching or exceeding the observed rate of polymorphism at CpG sites. Between 10.5 and 12.3% of SNPs occurred in CpG sites for each lake in contrast with an average of 3.8% of polymorphic sites across the genome. This means that 2.8 to 3.2 times more SNPs occur in CpG sites than in the rest of the genome. There was significant enrichment for all SNP types in CpG sites according to Pearson’s chi-squared tests (p < 0.001; Table 2). C/T and G/A SNPs were most common in CpG sites, with 3.0 to 3.4 times more C/T SNPs occurring in CpG sites than in all C and G sites across the genome. The other SNP types were slightly less enriched, with an enrichment of 1.1 to 1.2 times more C/G and G/C SNPs and 1.2 to 1.3 times more C/A and G/T SNPs in CpG sites relative to the rest of the C and G sites in the genome (Table 2).

**Table 2:** Pearson’s chi-squared test results for the rate of polymorphism in CpG sites (i) for all SNP types relative to the rate of polymorphism across the entire genome, and (ii) for specific SNP types relative to all C and G positions in the genome. Fold change indicates the ratio between observed and expected polymorphism rates.

| SNP Type | Value | Cliff | Indian | Langfjordvatn | Zurich |
| --- | --- | --- | --- | --- | --- |
| All | SNPs | 11 861 765 | 12 727 697 | 30 919 358 | 24 221 888 |
|  | SNPs in CpGs | 1 250 313 | 1 349 717 | 3 795 855 | 2 879 669 |
|  | Polymorphic CpGs (%) | 10.5 | 10.6 | 12.3 | 11.9 |
|  | Polymorphic sites (%) | 3.8 | 3.8 | 3.8 | 3.8 |
|  | Fold change | 2.8 | 2.8 | 3.2 | 3.1 |
|  | p-value (Chi-squared test) | < **0.001** | < **0.001** | < **0.001** | < **0.001** |
|  | p-value (permutation test) | < **0.001** | < **0.001** | < **0.001** | < **0.001** |
| C/T and G/A | SNPs | 2 804 547 | 3 080 961 | 8 321 890 | 6 251 941 |
|  | SNPs in CpGs | 750 311 | 819 314 | 2 487 283 | 1 834 278 |
|  | SNPs in CpGs (%) | 0.98 | 1.09 | 3.94 | 2.81 |
|  | SNPs in all Cs and Gs (%) | 0.32 | 0.36 | 1.17 | 0.85 |
|  | Fold change | 3.0 | 3.0 | 3.4 | 3.3 |
|  | p-value (Chi-squared test) | < **0.001** | < **0.001** | < **0.001** | < **0.001** |
| C/G and G/C | SNPs | 956 737 | 1 012 702 | 2 253 168 | 1 810 523 |
|  | SNPs in CpGs | 95 721 | 100 779 | 239 633 | 198 157 |
|  | SNPs in CpGs (%) | 0.13 | 0.13 | 0.38 | 0.30 |
|  | SNPs in all Cs and Gs (%) | 0.11 | 0.12 | 0.32 | 0.25 |
|  | Fold change | 1.1 | 1.1 | 1.2 | 1.2 |
|  | p-value (Chi-squared test) | < **0.001** | < **0.001** | < **0.001** | < **0.001** |
| C/A and G/T | SNPs | 2 124 638 | 2 314 058 | 5 422 615 | 4 126 260 |
|  | SNPs in CpGs | 234 886 | 248 499 | 590 917 | 476 256 |
|  | SNPs in CpGs (%) | 0.31 | 0.33 | 0.94 | 0.73 |
|  | SNPs in all Cs and Gs (%) | 0.25 | 0.27 | 0.76 | 0.56 |
|  | Fold change | 1.3 | 1.2 | 1.2 | 1.3 |
|  | p-value (Chi-squared test) | < **0.001** | < **0.001** | < **0.001** | < **0.001** |

### Epigenetic differentiation between benthic and limnetic whitefish

We assessed the level of epigenetic differentiation between benthic-limnetic species pairs by performing differential methylation analysis for each lake. After all quality trimming, we analyzed between 12 449 354 and 18 999 942 CpG sites per lake with approximately 7.5 to 8.1X coverage per sample (see Table 3 for detailed information). We identified variable but significant epigenetic differentiation between benthic-limnetic species pairs in all lakes: 38 060 differentially methylated loci (DMLs, i.e., CpG sites) and 2 891 differentially methylated regions (DMRs, i.e., prolonged regions of the DNA with differences in methylation between species) in Cliff Lake, 24 949 DMLs and 2 300 DMRs in Indian Lake, 3 537 DMLs and 367 DMRs in Langfjordvatn Lake, and 7 140 DMLs and 703 DMRs in Zurich Lake (Figure 2, Table 3, Tables S1-S8). The DMRs covered between 2 296 and 24 524 CpG sites in each lake (Table 3). Overall epigenetic differentiation between species was greater in North America than in Europe, consistent with higher interspecies genetic differentiation as measured by F_ST_ in North America relative to Europe.

**Fig. 2:**
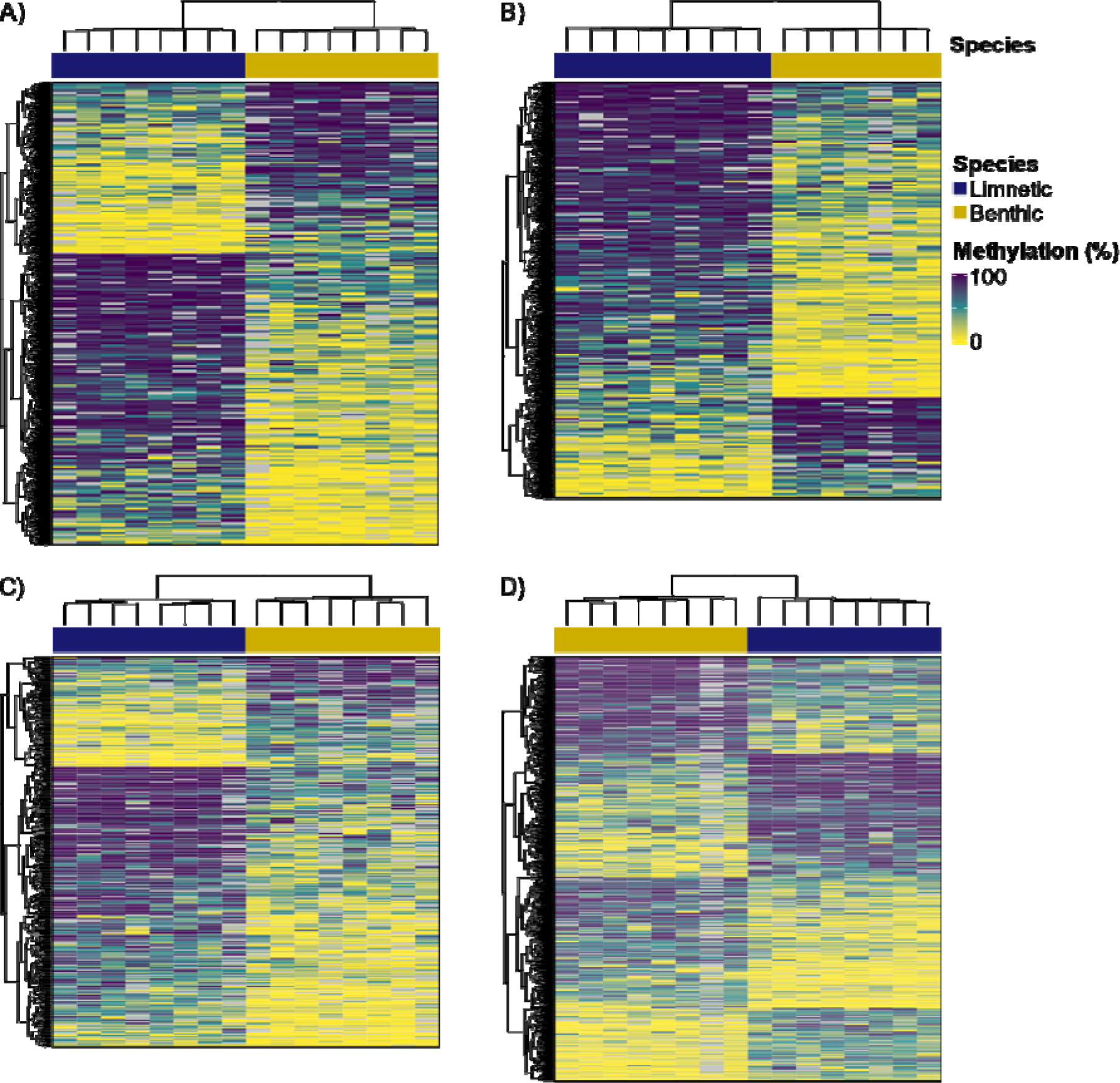
Differential methylation analysis comparing limnetic-benthic species pairs identified (A) 2 891 DMRs in Cliff Lake, (B) 2 300 DMRs in Indian Lake, (C) 367 DMRs in Langfjordvatn Lake, and (D) 703 DMRs in Zurich Lake. Limnetic (navy) and benthic (gold) species are generally neatly differentiated in the dendrograms based on Euclidean distance shown on the x-axes. Percent methylation is displayed for each DMR from 0% (yellow) to 100% (indigo) with clear DNA methylation differences between species. Each column represents a sample and each row represents a DMR.

**Table 3:** Statistics from differential methylation analysis, including coverage depth and number of CpGs analysed after all coverage trimming, number of DMLs and DMRs identified using a significance level of p < 0.05, and information on outlier SNP-DML overlaps and Pearson chi-squared test results.

| Statistic | Cliff | Indian | Langfjordvatn | Zurich |
| --- | --- | --- | --- | --- |
| Average coverage depth (WGBS) | 7.5 | 7.6 | 8.1 | 8.1 |
| Total CpGs analyzed | 16 668 063 | 18 999 942 | 12 449 354 | 17 896 958 |
| DMLs | 38 060 | 24 949 | 3 537 | 7 140 |
| % CpGs that are DMLs | 0.2 | 0.1 | 0.03 | 0.04 |
| DMRs | 2891 | 2300 | 367 | 703 |
| CpGs in DMRs | 24524 | 19566 | 2296 | 4547 |
| %CpGs in DMRs | 0.15 | 0.10 | 0.02 | 0.03 |
| Outlier SNPs (5% highest $F_{ST}$) | 593088 | 636384 | 1545963 | 1211091 |
| Outlier SNP-DML overlaps | 180 | 178 | 45 | 132 |
| Expected frequency of overlaps | 1.38e-05 | 2.62e-05 | 9.12e-06 | 1.30e-05 |
| Observed frequency of overlaps | 9.08e-04 | 7.71e-04 | 7.65e-05 | 2.07e-04 |
| Fold enrichment | 65.9 | 29.4 | 8.4 | 15.9 |
| p-value | **<0.001** | **<0.001** | **<0.001** | **<0.001** |

### Enrichment for overlaps between outlier SNPs and DMLs

For each lake, we assessed whether the most differentiated SNPs (5% highest F_ST_) in each lake overlap more often than expected by chance with CpG sites identified as DMLs between species. Such overlaps in genetic and epigenetic differentiation could represent potential sites of ongoing genetic divergence where divergent methylation patterns could be undergoing epigenetically influenced mutagenesis into stable genetic variants between species. We considered only polymorphic CpG sites in this analysis to account for the elevated mutation rate at CpG sites. Pearson’s chi-squared tests showed that the number of outlier SNP-DML overlaps was much greater than expected based on the combined probability of finding both an outlier SNP and a DML in a polymorphic CpG site (Table 3; see methods). We found between an 8.4 to a 65.9-fold enrichment of overlaps depending on lake. The enrichment was greater in North America (65.9-fold in Cliff Lake and 29.4-fold in Indian Lake) than in Europe (8.4-fold in Langfjordvatn Lake and 15.9-fold in Zurich Lake).

Gene ontology enrichment analysis for the outlier SNP-DML overlaps showed enrichment for 12 terms mostly involved in immune function in Cliff Lake, no terms in Indian Lake, five molecular function terms in Langfjordvatn Lake, and 10 terms in Zurich Lake associated with DNA binding and transcription (Table 4).

**Table 4:** Gene ontology results for outlier SNP-DML overlaps in lake and European whitefish. There were 12 enriched terms from Cliff Lake, none from Indian Lake, five from Langfjordvatn Lake, and 10 from Zurich Lake. Significant results had a Benjamini-Hochberg false discovery rate corrected p-value less than 0.1 and were filtered for GO term depth greater than 1 to remove broad terms.

| GO term | Subontology | GO term name | GO term depth | p-value (BH-FDR) |
| --- | --- | --- | --- | --- |
| **Cliff Lake** |  |  |  |  |
| GO:0050912 | BP | detection of chemical stimulus involved in sensory perception of taste | 5 | 0.047 |
| GO:0002414 | BP | immunoglobulin transcytosis in epithelial cells | 6 | 0.044 |
| GO:0001580 | BP | detection of chemical stimulus involved in sensory perception of bitter taste | 6 | 0.047 |
| GO:0002415 | BP | immunoglobulin transcytosis in epithelial cells mediated by polymeric immunoglobulin receptor | 7 | 0.044 |
| GO:0071745 | CC | IgA immunoglobulin complex | 3 | 0.003 |
| GO:0042571 | CC | immunoglobulin complex, circulating | 3 | 0.009 |
| GO:0071746 | CC | IgA immunoglobulin complex, circulating | 4 | 0.003 |
| GO:0071749 | CC | polymeric IgA immunoglobulin complex | 5 | 0.003 |
| GO:0071751 | CC | secretory IgA immunoglobulin complex | 6 | 0.003 |
| GO:0019763 | MF | immunoglobulin receptor activity | 4 | 0.042 |
| GO:0005154 | MF | epidermal growth factor receptor binding | 5 | 0.013 |
| GO:0001792 | MF | polymeric immunoglobulin receptor activity | 5 | 0.013 |
| **Langfjordvatn Lake** |  |  |  |  |
| GO:0016840 | MF | carbon-nitrogen lyase activity | 3 | 0.052 |
| GO:0016842 | MF | amidine-lyase activity | 4 | 0.014 |
| GO:0016715 | MF | oxidoreductase activity, acting on paired donors, with incorporation or reduction of molecular oxygen, reduced ascorbate as one donor, and incorporation of one atom of oxygen | 4 | 0.015 |
| GO:0004504 | MF | peptidylglycine monooxygenase activity | 5 | 0.001 |
| GO:0004598 | MF | peptidylamidoglycolate lyase activity | 5 | 0.001 |
| **Zurich Lake** |  |  |  |  |
| GO:0003700 | MF | DNA-binding transcription factor activity | 2 | 0.027 |
| GO:0000981 | MF | DNA-binding transcription factor activity, RNA polymerase II-specific | 3 | 0.030 |
| GO:0001067 | MF | transcription regulatory region nucleic acid binding | 4 | 0.051 |
| GO:0043565 | MF | sequence-specific DNA binding | 5 | 0.051 |
| GO:0003690 | MF | double-stranded DNA binding | 5 | 0.090 |
| GO:1990837 | MF | sequence-specific double-stranded DNA binding | 6 | 0.061 |
| GO:0000976 | MF | transcription cis-regulatory region binding | 7 | 0.051 |
| GO:0000987 | MF | cis-regulatory region sequence-specific<br>DNA binding | 8 | 0.010 |
| GO:0000977 | MF | RNA polymerase II transcription regulatory<br>region sequence-specific DNA binding | 8 | 0.030 |
| GO:0000978 | MF | RNA polymerase II cis-regulatory region<br>sequence-specific DNA binding | 9 | 0.010 |

### Gene-level parallelism for epigenetic variation among lakes

We assessed parallelism in the previously identified DMLs, DMRs, and outlier SNP-DML overlaps between benthic and limnetic species among lakes. We were unable to compare exact genomic locations for these markers since we used different reference genomes for European and lake whitefish. Instead, we determined which markers directly overlapped with gene transcripts in the annotated whitefish genomes and considered parallelism in genes across lakes (Figure 3). There was some parallelism in the genes associated with DMLs and DMRs (111 and three common genes among all four lakes representing 2.0% and 0.23% of genes, respectively), especially shared between two or three lakes. There was little parallelism with respect to outlier SNP-DML overlaps with common genes among all lakes.

**Fig. 3:**
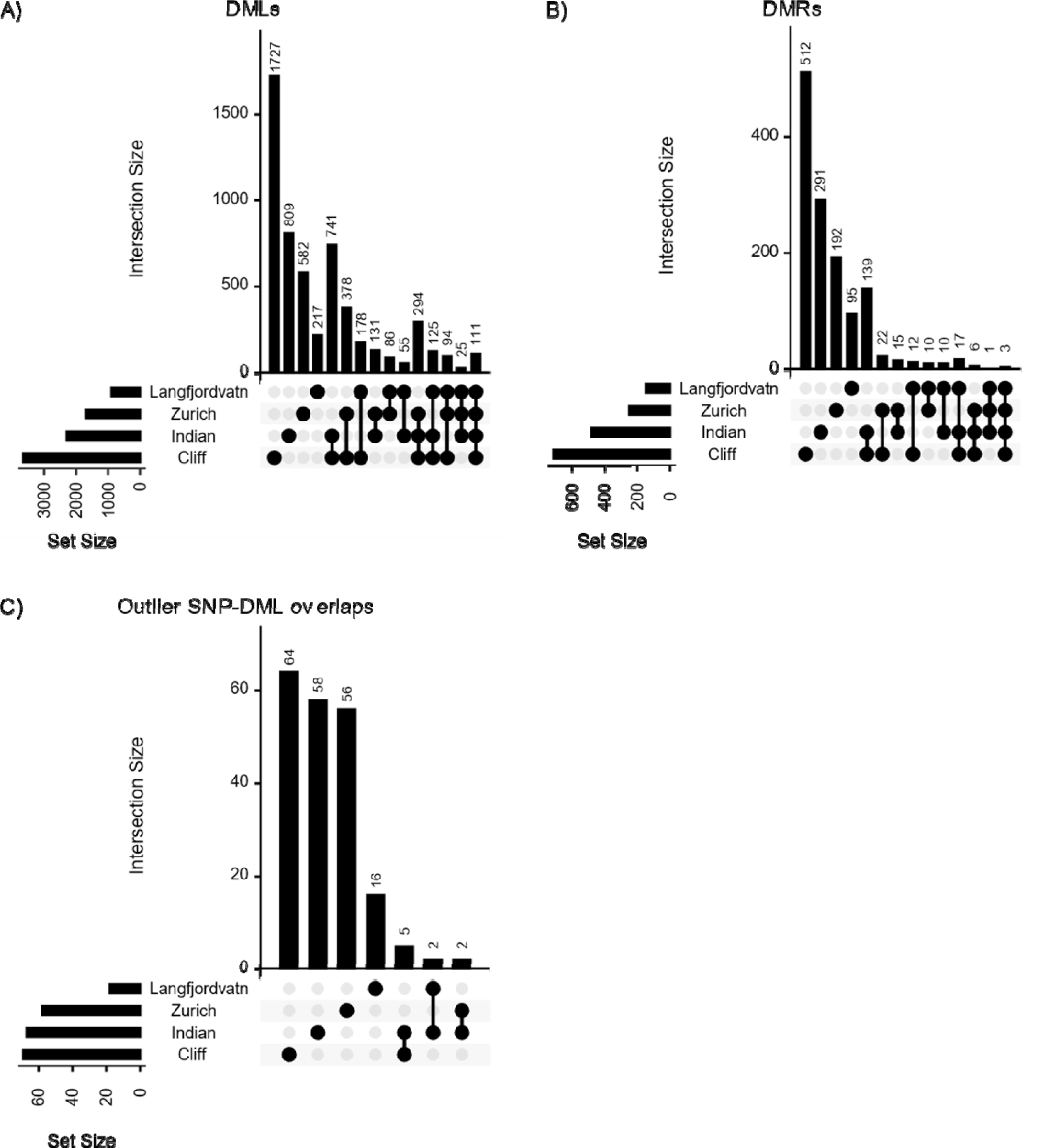
Parallelism among lakes was assessed at the gene level based on the number of unique genes overlapping (A) DMLs, (B) DMRs, and (C) outlier SNP-DML overlaps. Missing comparisons indicate that no genes are shared between those lakes. The connected dots on the x-axis represent the populations considered in each comparison and the intersection size shows the number of parallel genes for each comparison. Set size gives the number of unique genes overlapping the marker in each lake.

## Discussion

Recent speciation research has expanded to include epigenetic mechanisms such as DNA methylation (Pál & Miklós 1999; Richards et al. 2010; Greenspoon et al. 2022; Ashe et al. 2021) which could contribute to phenotypic diversification and reproductive isolation (Laporte et al. 2019). Here we provide evidence for genetic and epigenetic differentiation between sympatric benthic-limnetic whitefish species pairs from two continents with limited epigenetic parallelism (i.e., shared genes) between lakes. We show that polymorphism is enriched at CpG sites, including high overlap of outlier SNPs and CpG sites showing differential methylation between species which may represent sites of ongoing mutagenesis. Together, our results provide support for the proposed contributions of DNA methylation to phenotypic diversification (Anastasiadi et al. 2021; Vogt 2021), mutagenesis (Tomkova & Schuster-Böckler 2018), and speciation (Pál & Miklós 1999; Richards et al. 2010; Greenspoon et al. 2022; Ashe et al. 2021).

### Considerable but variable genetic divergence between species and lakes

We observed genetic differentiation between species in all four lakes, consistent with previous studies (Rougeux et al. 2017; Bernatchez et al. 2010; Dion-Côté et al. 2017; Rogers & Bernatchez 2007; Rougeux, Gagnaire, Praebel, et al. 2019; Østbye, Næsje, et al. 2005; Østbye et al. 2006; Siwertsson et al. 2013; De-Kayne et al. 2022; Feulner & Seehausen 2019; Mérot et al. 2023). There was greater genetic differentiation between species in Cliff Lake than Indian Lake in North America, as previously reported in our sister study which reported on the same lake whitefish data and conclusions (Mérot et al. 2023) and in another study with lower genomic resolution (Gagnaire et al. 2013). Genetic differentiation was greater between lakes than between species in Europe (Table 1B), likely due to the Zurich Lake and Langfjordvatn Lake populations having occupied two different glacial refugia during the last glaciation period (Østbye, Bernatchez, et al. 2005). However, our within-lake genome-wide F_ST_ estimates are slightly low compared to other estimates between European whitefish species pairs (0.037 to 0.12 based on RAD sequencing (Feulner & Seehausen 2019), 0.042 to 0.096 based on 16 microsatellite loci (Siwertsson et al. 2013) and 0.01 to 0.075 based on six microsatellite loci (Østbye et al. 2006)), likely due to the use of whole genome vs. targeted or reduced representation methods and the low MAF filter (MAF > 0.05) including many low frequency SNPs in this analysis. Variable interspecies F_ST_ between lakes is consistent with varying degrees of reproductive isolation/gene flow between species in different systems (Lu & Bernatchez 1999; Gagnaire et al. 2013; De-Kayne et al. 2022; Rougeux, Gagnaire & Bernatchez 2019; Østbye et al. 2006). They are also consistent with estimated lake whitefish divergence history in Cliff and Indian Lakes (32 000 years in allopatric speciation and 9 200 years since secondary contact for Cliff, 29 000 years in allopatric speciation and 8 500 years since secondary contact for Indian) (Rougeux et al. 2017). European whitefish divergence is estimated to have occurred over a longer timeline (121 000 years in allopatry and 12 000 years since secondary contact for Langfjordvatn, 107 000 years in allopatry and 28 600 years since secondary contact for Zurich) (Rougeux, Gagnaire & Bernatchez 2019). Despite European whitefish having a longer period of allopatric speciation, lake whitefish show greater genetic divergence. The variation in interspecies genetic divergence between lakes could be due to the extent of differences between the species’ trophic niches, with species occupying vastly different trophic niches also showing greater morphological differences (Lu & Bernatchez 1999). Overall, genetic differentiation between species was widespread along the genome consistent with previous work (De-Kayne et al. 2022; Feulner & Seehausen 2019; Mérot et al. 2023), though large-effect loci (De-Kayne et al. 2022) and genomic islands of divergence (Gagnaire et al. 2013) have also been reported. This genome-wide genetic differentiation between species may be associated with phenotypic divergence and reproductive isolation between species (Coyne & Orr 2004).

### Epigenetic divergence between whitefish species pairs

We found considerable epigenetic divergence in liver tissue between species, though the extent of divergence varied between lakes and was greater in North America than in Europe (Figure 2, Table 3). Our results support the idea that DNA methylation could provide an additional mechanism for phenotypic diversification and speciation, similar to results in lake whitefish showing differential methylation of transposable elements between species in liver tissue, which can affect transposable element activity (Laporte et al. 2019). These methylation changes may have arisen due to environmental differences between benthic and limnetic habitats which could contribute to character displacement between species since the environment can have profound effects on the methylome. Previous studies have shown that methylation is affected by temperature (Ryu et al. 2018; Venney et al. 2022; Metzger & Schulte 2017; McCaw et al. 2020; Anastasiadi et al. 2017), salinity (Heckwolf et al. 2020; Artemov et al. 2017), rearing environment (Leitwein et al. 2021; Le Luyer et al. 2017; Wellband et al. 2021; Gavery et al. 2018), and other factors in various systems. Differences in habitat use can influence DNA methylation, as observed between capelin (*Mallotus villosus*) utilizing beach- and demersal-spawning life history tactics (Venney et al. 2023), among sympatric Arctic charr morphs occupying different habitats and dietary niches (Matlosz et al. 2022), between phenotypically divergent cichlid species (Vernaz et al. 2022), and in freshwater snails (*Potamopyrgus antipodarum*) which exhibit differences in shell shape associated with water current speed (Thorson et al. 2017).

DNA methylation can also be influenced by genetic variation (Richards 2006; Lallias et al. 2021), therefore epigenetic differences may in part be driven by genetic divergence between species. Given the greater epigenetic divergence and F_ST_ between species pairs in North America than in Europe, it is possible that the methylation differences are a function of genetic divergence between species. Parallel transcriptional differences were previously reported between benthic-limnetic whitefish species pairs in both captive and natural conditions, indicating that some transcriptional differences between species are under genetic control and may have been subject to selection (St-Cyr et al. 2008). Transcriptional divergence between benthic-limnetic species pairs was also shown in 48 of 64 samples used in this study (6 samples per species per lake) and differences were often parallel across lakes in both lake whitefish and European whitefish (see Rougeux, Gagnaire, Praebel, et al. 2019). We generally observed similar ratios of interspecies DMRs among lakes from our study and differentially expressed gene (DEG) counts in the earlier transcriptome study by Rougeux et al. (2 891 DMRs and 3 175 DEGs in Cliff Lake, 2 300 DMRs and 238 DEGs in Indian Lake, 367 DMRs and 276 DEGs in Langfjordvatn Lake, and 703 DMRs and 1 392 DEGs in Zurich Lake). This indicates that Cliff Lake and Zurich Lake have greater epigenomic and transcriptomic differences between species relative to the other lakes, though there are considerably more DMRs than DEGs in Indian Lake, possibly due to greater genetic and epigenetic differentiation between species. Thus, DNA methylation divergence between species may be due to differences in habitat and genetic background and could provide a mechanistic basis for phenotypic diversification between these nascent species pairs due to its effects on transcription (Bird 2002) and phenotype (Anastasiadi et al. 2021; Vogt 2021).

### Mutational enrichment at CpG sites supports epigenetically influenced mutagenesis

Our results suggest that polymorphism is enriched at CpG sites, possibly due to the mutagenic nature of DNA methylation (Flores et al. 2013; Tomkova & Schuster-Böckler 2018; Ashe et al. 2021) generating genetic variation between species pairs. We show that CpG sites, the main sites where DNA methylation occurs in vertebrate genomes, have higher levels of polymorphism compared to the rest of the genome (Table 2). This suggests that DNA methylation may be inducing point mutations that could accumulate between species over generations. We observed a 2.8- to 3.2-fold polymorphism enrichment at CpG sites which is comparable to the previously reported 3.5-fold increased mutation in methylated relative to unmethylated cytosines (Gorelick 2003; Jones et al. 1992). It is likely that more mutations have occurred yet were not detected due to selection against them since epigenetically induced mutations are generally inferred to be deleterious based on mismatches between experimentally observed epigenetically induced mutation rate and the observed rates of these mutations in populations (Danchin et al. 2019). However, some mutations could be retained through random genetic drift if neutral or mildly deleterious, or selected for if they prove beneficial, leading to increased frequency of the novel mutation over time. While epigenetically induced C>T transitions are most common due to spontaneous deamination of cytosines to uracil (Tomkova & Schuster-Böckler 2018), we show that there is significant enrichment of all types of point mutations at CpG sites (Table 2). The exact mechanisms behind spontaneous C>A and C>G mutations without the involvement of mutagens are less clear in the context of epigenetically influenced mutagenesis (Tomkova & Schuster-Böckler 2018), though we provide evidence that all types of polymorphism are enriched at CpG sites.

### Co-occurrence of genetic and epigenetic divergence at CpG sites

Our results showed that SNPs with high F_ST_ between species are enriched at DMLs. These sites could potentially reflect ongoing mutagenesis or genetic assimilation contributing to genetic evolution between benthic-limnetic species pairs, wherein one species is in the process of losing the CpG site. While the ancestral species is unknown, it is also possible that some of these sites have mutated from a non-CpG site to a CpG site (e.g., from TpG to CpG), though a causative mechanism for this is not immediately clear. The outlier SNP-DML overlaps showed both differential liver methylation between species (i.e., when CpG sites are still present in both species) and high genetic differentiation between species (i.e., when one species has partially assumed the “assimilated” state and a cytosine has mutated to another nucleotide). There was a greater enrichment for outlier SNP-DML overlaps in North America than Europe (see Table 3) consistent with greater genetic differentiation and more pronounced reproductive isolation between species pairs in North America, though this could also reflect differences in mutational signatures between species (e.g., Goldberg and Harris 2022). It is also likely that the degree of enrichment would differ among tissues and through ontogeny since the methylome is tissue-specific (Venney et al. 2016; Christensen et al. 2009; Gavery et al. 2018) and affected by age (Venney et al. 2016; Christensen et al. 2009). Gene ontology analysis of the overlaps showed that they occurred in genes associated with behaviour, immune function, metabolism, DNA binding, cellular processes, oxio-reductase activity, and transcription factor activity depending on the lake (Table 4), though there were no shared GO terms among lakes in our study. Nevertheless, our findings were consistent with previous transcriptomic studies in whitefish showing enrichment of similar functions in DEGs (Rougeux, Gagnaire, Praebel, et al. 2019; St-Cyr et al. 2008), with potential implications for immune function and growth (Rougeux, Gagnaire, Praebel, et al. 2019). To our knowledge, this one of the first studies to relate epigenetic divergence and putatively epigenetically induced polymorphism to speciation, providing support for previous theories on the role of DNA methylation in genetic assimilation. Stajic et al. (2019) previously showed that mutational assimilation was dependent on the capacity of *S. cerevisiae* to modify histone acetylation, showing the involvement of epigenetic mechanisms in genetic assimilation. A recent study in threespine stickleback (*Gasterosteus aculeatus*) showed that DMLs between freshwater and marine stickleback were also associated with high nucleotide diversity (Ord et al. 2023). Interestingly, a study in great apes (*Homo*, *Pan*, *Gorilla*, and *Pongo* sp.) found that mutational signatures across the genome differed between species and that epigenetically induced mutations were influenced by chromatin state and cytosine hydroxymethylation (Goldberg & Harris 2022). The role of epigenetic processes in inducing mutagenesis is becoming clearer, though we provide some initial support for DNA methylation influencing mutagenesis in the context of ecological speciation.

While the idea of genetic assimilation is exciting, there are other explanations for the association between outlier SNPs and DMLs. An alternative hypothesis is that both DNA methylation and genetic polymorphism at the same CpG site could have similar effects on transcription and thus provide two different, simultaneous molecular mechanisms for controlling transcription at the same site in different individuals. Genetic variation at proximal linked sites could also determine methylation state at these genetically and epigenetically differentiated CpG sites, though it is unclear why this would cause an enrichment of outlier SNP-DML overlaps given widespread genomic differentiation between species in all lakes. It is also possible that selection maintains both genetic and epigenetic state at these sites and the two are not related. Future long-term experimental evolution studies have the potential to distinguish between these possibilities and facilitate real-time observation of epigenetically induced mutagenesis and genetic assimilation.

### A hypothetical role for DNA methylation in whitefish speciation

When environments change and then remain stable (e.g., in range expansions, habitat colonization, or shifts to novel habitat use), DNA methylation may serve as an initial plastic response to the new environment, with the new methylation state being maintained by natural selection (Ashe et al. 2021). Given the rapid postglacial speciation rate (∼3-4K generations) between benthic and limnetic species in all lakes, methylation differences could have arisen quickly, contributing to the multi-trait rapid evolution that occurred between species. Over time, epigenetically influenced mutagenesis could have led to methylation changes inducing genetic divergence (Danchin et al. 2019; Ashe et al. 2021). Therefore, epigenetically induced mutagenesis could have contributed to genetic differentiation between benthic and limnetic whitefish. Divergent selection could then have acted on both epigenetic and genetic marks if they affected phenotype. This is consistent with heritable differences in behaviour (e.g., Rogers et al. 2002) and minimal plasticity in morphological traits differentiating benthic and limnetic lake whitefish (Laporte et al. 2016). Transcriptomic differences between benthic-limnetic species pairs were also stable across environments in lake whitefish (St-Cyr et al. 2008) and related to parallel genetic differences between species in both lake and European whitefish (Rougeux, Gagnaire, Praebel, et al. 2019), suggesting that these putatively adaptive traits might be genetically controlled. However, further study would be needed for any direct test of environmental differences between species pairs or a link between (epi)genetic variation and phenotype.

### Conclusions

We found substantial genetic and epigenetic divergence between independently derived benthic-limnetic species pairs in lake and European whitefish. We provide evidence that DNA methylation may lead to increased polymorphism due to its mutagenic nature, potentially contributing to early phenotypic diversification, genetic divergence, and speciation. We characterized potential sites of ongoing genetic assimilation wherein differential methylation levels between species may be influencing polymorphism and resulting in divergent genetic variation between benthic-limnetic species pairs. As such, our results shed light on the diverse ways DNA methylation can contribute to plasticity and evolution, from plastic responses to environmental changes to the induction of mutagenesis leading to genetic divergence. Future studies using experimental evolution or evolve and resequence approaches are needed to confirm the mutagenic nature of DNA methylation and its role in genetic assimilation.

## Materials and Methods

### Sample preparation and sequencing

We sampled limnetic-benthic whitefish species pairs from two lakes in North America and two lakes in Europe: lake whitefish (*C. clupeaformis*) from Cliff and Indian Lake, Maine, USA, and European whitefish (*C. lavaretus*) from Langfjordvatn Lake, Norway and Zurich Lake, Switzerland (Fig. 1). Animal care was performed humanely under Université Laval animal care permit 126316. Fish were caught with gillnets, humanely euthanized, and immediately dissected to obtain fresh tissue samples. Liver tissue was sampled from 64 fish, including six per lake previously used in Rougeux et al. (2019): eight individuals per species per lake except for Indian Lake where we sampled seven benthic and nine limnetic fish. Samples were stored either at -80°C or in RNAlater; all samples from Europe were stored in RNAlater. Liver tissue was chosen due to its homogeneous tissue characteristics and involvement in growth and metabolism (Trefts et al. 2017).

Genomic DNA was isolated using a modified salt extraction protocol (Aljanabi & Martinez 1997). DNA was checked on a 1% agarose gel and quantified on a NanoDrop spectrophotometer. WGS and WGBS libraries were built at the McGill University and Genome Quebec Innovation Centre (Montreal, Canada) using in-house protocols. WGS was performed using paired end 150 bp sequencing on an Illumina HiSeq4000 with estimated 5X coverage. The North American samples were previously sequenced in Mérot et al. (2022), and WGBS was performed in this study only using paired end 150 bp sequencing on the Illumina HiSeqX with samples randomly distributed across 16 lanes (four per lane) with ∼10X coverage.

### Whole genome sequencing analysis

Reads were trimmed and quality filtered with fastp (Chen et al. 2018). The North American samples were aligned to the *Coregonus clupeaformis* genome (ASM1839867v1; (Mérot et al. 2023) and the European samples were aligned to the European whitefish genome (*Coregonus* sp. Balchen; LR778253.1; De-Kayne et al. 2020) using BWA-MEM (Li 2021). Aligned reads were filtered to require mapping quality over 10 with Samtools v1.8 (Li et al. 2009). Duplicate reads were removed with MarkDuplicates (PicardTools v1.119., http://broadinstitute.github.io/picard). We realigned around indels with GATK IndelRealigner (McKenna et al. 2010) and soft clipped overlapping read ends using clipOverlap in bamUtil v1.0.14 (Breese & Liu 2013). The pipeline is available at https://github.com/enormandeau/wgs_sample_preparation.

Bam alignments were analysed with the program ANGSD v0.931 (Korneliussen et al. 2014) which accounts for genotype uncertainty and is appropriate for low and medium coverage WGS (Lou et al. 2021). Reads were filtered to remove low-quality reads and to keep mapping quality above 30 and base quality above 20. We ran ANGSD on each lake separately to detect polymorphic positions (SNPs), estimate the spectrum of allele frequency, minor allele frequency (MAF), and genotype likelihoods. Genotype likelihoods were estimated with the GATK method (-GL 2). The major allele was the most frequent allele (-doMajorMinor 1). We kept positions covered by at least one read in at least 75% of individuals, with a total coverage below 400 (25 times the number of individuals) to avoid including repeated regions in the analysis. After such filtering, we exported a list of covered positions for each lake for further analysis, including 1 998 994 058 positions in Cliff Lake, 1 981 247 030 in Indian Lake, 1 658 793 591 in Langfjordvatn, and 1 702 852 520 in Zurich Lake.

From this list of variant and invariant positions, we extracted a list of SNPs as the variable positions with an MAF above 5% and subsequently used this list with their respective major and minor alleles for most analyses (i.e., F_ST_ between benthic and limnetic, overlap with CpG sites). Differentiation between benthic and limnetic species in each lake was measured with F_ST_ statistics, using ANGSD to estimate joint allele frequency spectrum, realSFS functions to compute F_ST_ in sliding windows of 100 KB with a step of 25 KB. Positions were restricted to the polymorphic SNPs (>5% MAF) previously polarized as major or minor allele (options –sites and –doMajorMinor 3), and which were covered in at least 75% of the samples in each species. F_ST_ estimates for Cliff Lake and Indian Lake in North America were previously published in Mérot et al. (2022). F_ST_ estimates between lakes were performed with the same analyses at the continent level to ensure a consistent polarisation of the major allele. The ANGSD pipeline is available at https://github.com/clairemerot/angsd_pipeline.

### Frequency of SNPs in CpG sites

We determined whether CpG sites were more polymorphic than the rest of the genome, which would be indicative of CpG methylation influencing mutagenesis (Jones et al. 1992; Gorelick 2003). We used fastaRegexFinder.py (https://github.com/dariober/bioinformatics-cafe/blob/master/fastaRegexFinder/) to find all CpG sites in each reference genome. Next, we restricted the CpG site lists to include only sites with sufficient coverage in the WGS data using bedtools *intersect* (Quinlan & Hall 2010) with the list of covered positions for each lake exported earlier (>75% of samples covered). We used regioneR (Gel et al. 2016) to determine how many SNPs fell within CpG sites using the *numOverlaps* command. We assumed a uniform distribution of SNPs across the genome as a null hypothesis for both tests, thus a higher proportion of SNPs in CpGs relative to average polymorphism in the genome would indicate an enrichment of point mutations in CpGs.

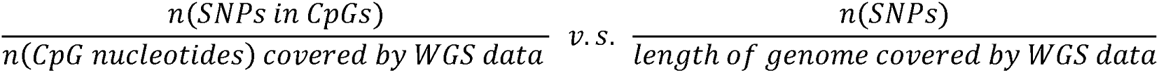

We also used a permutation test to determine if the enrichment of SNPs in CpG sites would occur by chance over 10 000 iterations. We determined the proportion of the genome made up of CpG sites by dividing the number of CpG nucleotides by the total length of the genome with sufficient WGS coverage. We assumed a uniform distribution of SNPs across the genome as a null hypothesis for both tests, thus a higher proportion of SNPs in CpG sites relative to the proportion of the genome made up of CpG sites would indicate an enrichment of point mutations in CpGs.

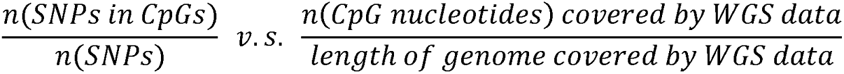

We tested whether there was an enrichment of specific SNP substitution types as C/T polymorphisms are expected to be more common due to deamination of methylated cytosines (Tomkova & Schuster-Böckler 2018; Gorelick 2003; Jones et al. 1992). For this analysis, we only considered C and G sites in the genome due to the increased mutagenicity of these nucleotides (Kiktev et al. 2018). We split the list of SNPs into three separate files for each possible mutation: (i) all C/T and G/A SNPs (where G/A SNPs would indicate a C/T mutation at the G position in the reverse complement of the DNA), (ii) all C/A and T/G SNPs, and (iii) all C/G SNPs. We also generated a list of all C and G sites in the genome with sufficient WGS coverage. We compared the rate of finding each SNP type in a CpG site to the rate of finding that SNP type in C and G sites, then used Pearson’s chi-squared test to determine if the proportions were significantly different. All scripts are available at https://github.com/cvenney/ga_permutation.

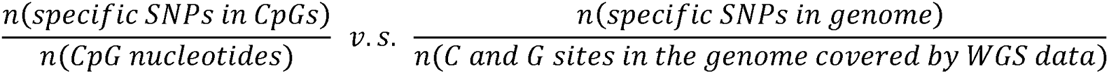

### Whole genome DNA methylation tabulation

Raw methylation data were trimmed using fastp (Chen et al. 2018) to remove sequences with phred quality less than 25, length less than 100 bp, and to remove the first and last nucleotides which have high sequencing error rate. After trimming, lake and European whitefish methylation data were analysed separately. Trimmed sequences were aligned to the same reference genomes as described in the WGS methods using bwa-meth (https://github.com/brentp/bwa-meth). Alignments with mapping quality greater than 10 were outputted to a BAM file using samtools (Li et al. 2009) and duplicate sequences were removed using Picard tools. Methyldackel’s *mbias* function was used to inform trimming of biased methylation calls at the beginning and end of reads (https://github.com/dpryan79/MethylDackel). CpG-specific methylation calling was performed on bias-trimmed data using methyldackel’s *extract* function while removing any detected variant sites where SNPs could affect methylation calling in that individual (*--maxVariantFrac 0.1 --minOppositeDepth 1*). The pipeline is available at https://github.com/enormandeau/bwa-meth_pipeline.

### Individual-level whitelisting of methylation data using SNP data

C/T SNPs cannot be discerned from true methylation reads because bisulfite conversion leaves methylated cytosine unchanged but converts unmethylated cytosine to uracil which is sequenced as a thymine. Therefore, CpG sites which overlap with C/T or G/A SNPs must be removed from the analysis. We used an individual-level approach where we filtered CpG sites separately for each sample based on that sample’s genotype (Figure 1), in addition to methylDackel’s built-in *--maxVariantFrac* SNP removal function described above. Therefore, we filter problematic SNPs out of this dataset using both the SNP and the methylation data. Relying on our WGS data, we thus created a whitelist of CpG sites for each sample, keeping the sites homozygous for C and G and masking individuals heterozygous or homozygous for the T or A allele at that site. To do so, we kept CpG sites that were covered in at least 12 individuals in a given lake (75% of samples for each lake). Then we filtered the whitelists, requiring sites to be either (i) non-variant positions, (ii) not a C/T SNP for the C position of the CpG in the lake, (iii) not a G/A SNP for the G position of the CpG in the lake, or (iv) a C/T SNP or a G/A SNP in other individuals within the lake, but where this individual has a likelihood greater than 0.7 to be a C/C and G/G homozygote. This resulted in individual-level SNP masking for each sample where CpG sites with C/T and G/A SNPs were removed in heterozygous individuals, but not blindly across all samples (see Figure 1 for a visual representation). C/C homozygotes and individuals with C/A and C/G SNPs at a given CpG site were retained in the analysis as these genotypes do not affect methylation calling. BedGraph files were filtered to include only CpG sites covered in the whitelist (i.e., where both the C and G positions were whitelisted). The pipeline for SNP masking and subsequent methylation analysis is available at https://github.com/cvenney/ga_methyl.

### Coverage filtration and differential methylation analysis

Whitelisted bedGraph files were filtered to exclude CpG sites with less than five and more than 100 reads. The files were imported into R (R Core Team 2022) where we kept only CpG sites with sufficient coverage in at least four limnetic and four benthic samples in the lake. BedGraph files were reformatted for further analysis with DSS (Park & Wu 2016). We then assessed the level of epigenetic differentiation between benthic-limnetic species pairs by performing differential methylation analysis for each lake. Methylation data were smoothed over 500 bp regions using the built-in moving average algorithm in DSS to control for spatial correlation of methylation levels among proximal CpGs. We ran generalized linear models in DSS to identify differentially methylated loci (DMLs; i.e., CpG sites with significantly different methylation levels between limnetic and benthic species) and differentially methylated regions (DMRs; regions of the genome showing differences in methylation levels between experimental groups) between species. DMLs were considered significant if the false discovery rate (FDR) corrected p-value was less than 0.05. DMRs were identified in DSS as regions with many statistically significant DMLs with p-values less than 0.05.

### Overlap between DMLs and outlier SNPs

We assessed whether there was an enrichment of outlier SNP-DML overlaps compared to the number of overlaps expected by chance given the frequency of polymorphic DMLs and the frequency of outlier SNPS in CpG sites. Enrichment could indicate epigenetically influenced mutagenesis and genetic assimilation of methylation changes into stable genetic variants. We generated a list of highly differentiated SNPs for each lake, retaining only the top 5% of SNPs with the greatest F_ST_ between species, hereafter called “outlier SNPs”. DML test files were converted to bed format for input into bedtools, then *intersect* was used to determine the number of overlaps between outlier SNPs and DMLs. We used only CpG sites that were (i) covered by WGS data, (ii) covered by WGBS data, and (iii) polymorphic to account for the observed elevated mutation rate in CpG sites compared to the rest of the genome. We then used Pearson’s chi-squared test to determine if there was an enrichment of observed DML-outlier SNP overlaps in polymorphic CpG sites compared to the expected rate of finding overlaps.

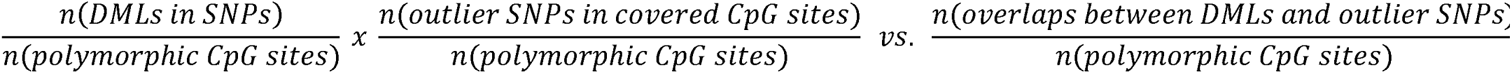

We also performed gene ontology (GO) enrichment analysis for the outlier SNP-DML overlaps for each lake. The whitefish genomes were first annotated with the GAWN pipeline (https://github.com/enormandeau/gawn) using the *Salvelinus namaycush* (GCF_018398675.1) transcriptome from GenBank, which is more complete than the available whitefish transcriptomes. We subsetted the annotation to include only the genes covered by both SNP and methylation data which represents the list of genes that could possibly contribute to enriched terms based on our data. We then retrieved the genes that overlapped the SNP-DML overlap positions for each lake and proceeded to GO enrichment tests using the pipeline at https://github.com/enormandeau/go_enrichment). Results were filtered to remove broad terms (depths 0 and 1) and to require a Benjamini-Hochberg FDR corrected p-value of 0.1 or less.

### Parallelism across lakes

We tested for parallelism in (epi)genetic variation among lakes for DMLs, DMRs, and outlier SNP-DML overlaps between benthic and limnetic species. Genomic positions of genes were identified using the annotated whitefish genomes from GO enrichment. Bedtools *intersect* was used to find direct overlaps between the gene positions and the markers of interest for each lake. Overlaps for each type of marker were visualized using UpSetR (Conway et al. 2017).

## Acknowledgements

We dedicate this article to the memory of Louis Bernatchez, who passed away in September 2023 during the review process. Louis had a tireless passion for science and whitefish, made outstanding contributions to the field, and was a wonderful mentor and friend. We thank Kim Præbel, Shripathi Bhat and the Freshwater ecology group at UiT (Tromso) for providing samples from Norway, and Ole Seehausen for providing samples from Switzerland. This work was supported by a Discovery research grant from the Natural Sciences and Engineering Research Council of Canada (NSERC) to L.B. CJV is supported by an NSERC postdoctoral fellowship.

## Data availability statement

European whitefish and lake whitefish whole genome sequencing data are available through SRA accessions PRJNA906116 and PRJNA820751, respectively. Whole genome bisulfite sequencing data are available through SRA accession PRJNA559821. All scripts are available on GitHub as indicated in the methods.

