## Supplementary material for "Epigenetic and genetic differentiation between *Coregonus* species pairs": Figure S1

### Epigenetic differentiation between *Coregonus* species pairs: potential for epigenetically induced mutation to contribute to speciation

Clare J Venney, Claire Mérot, Eric Normandeau, Clément Rougeux, Martin Laporte, Louis Bernatchez

**Figure S1:** Manhattan plots for Cliff Lake, Indian Lake, Langfjordvatn Lake, and Zurich Lake showing FST across the genome, calculated using 100 KB sliding windows with a step size of 25 KB. Chromosomes are plotted sequentially along the x-axis.

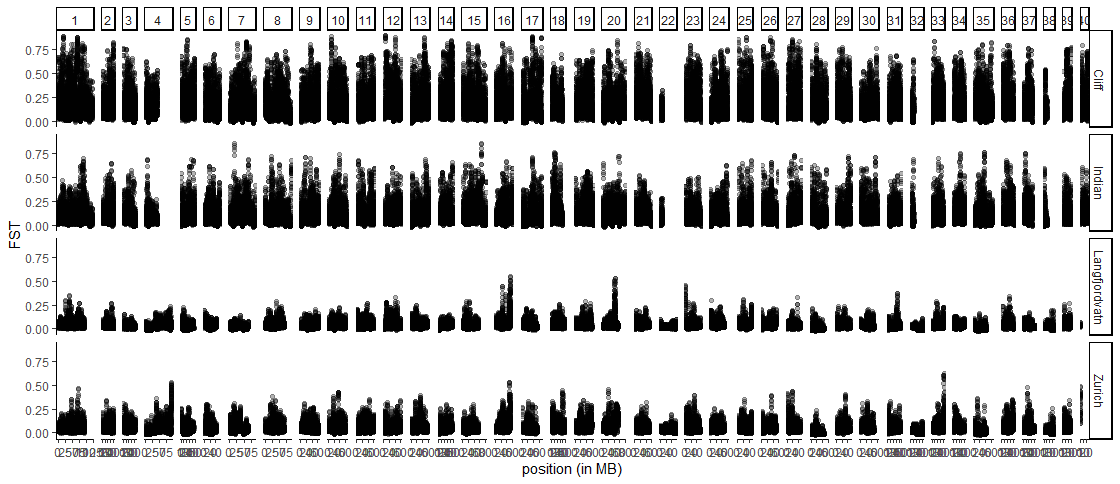
